# Occupancy distributions of membrane proteins in heterogeneous liposome populations

**DOI:** 10.1101/657098

**Authors:** Lucy Cliff, Rahul Chadda, Janice L. Robertson

**Affiliations:** Department of Molecular Physiology and Biophysics, The University of Iowa, Iowa City, IA; Department of Chemistry, The University of Bath, Bath, England, UK; Department of Biochemistry and Molecular Biophysics, Washington University, St. Louis, MO

**Keywords:** Liposome size distribution, Poisson, membrane protein, reconstitution, single-molecule photobleaching, CLC-ec1

## Abstract

Measurements of membrane protein structure and function often rely on reconstituting the protein into lipid bilayers through the formation of liposomes. Many measurements conducted in proteoliposomes, e.g. transport rates, single-molecule dynamics, monomer-oligomer equilibrium, require some understanding of the occupancy statistics of the liposome population for correct interpretation of the results. In homogenous liposomes, this is easy to calculate as the act of protein incorporation can be described by the Poisson distribution. However, in reality, liposomes are heterogeneous, which alters the statistics of occupancy in several ways. Here, we determine the liposome occupancy distribution for membrane protein reconstitution while taking into account liposome size heterogeneity. We calculate the protein occupancy for a homogenous population of liposomes with radius *r* = 200 nm, representing an idealization of vesicles extruded through 400 nm pores and compare it to the right-skewed distribution of 400 nm 2:1 POPE:POPG vesicles. As is the case for *E. coli* polar lipids, this synthetic composition yields a sub-population of small liposomes, ∼25 nm in radius with a long tail of larger vesicles. Previously published microscopy data of the co-localization of the CLC-ec1 Cl^-^/H^+^ transporter with liposomes, and vesicle occupancy measurements using functional transport assays, shows agreement with the heterogeneous 2:1 POPE:POPG population. Next, distributions of 100 nm and 30 nm extruded 2:1 POPE:POPG liposomes are measured by cryo-electron microscopy, demonstrating that extrusion through smaller pores does not shift the peak, but reduces polydispersity arising from large liposomes. Single-molecule photobleaching analysis of CLC-ec1-Cy5 shows the 30 nm extruded population increases the ‘Poisson-dilution’ range, reducing the probability of vesicles with more than one protein at higher protein/lipid densities. These results demonstrate that the occupancy distributions of membrane proteins into vesicles can be accurately predicted in heterogeneous populations with experimental knowledge of the liposome size distribution.

## 1. Introduction

Membrane protein reconstitution involves the incorporation of purified membrane-embedded proteins into lipid bilayers for functional and structural interrogation (1). It provides a means to study the protein in the context of the native solvent structure, which is a distinctly different environment than the detergent micelle state from which the protein is usually purified. There are several methods of reconstituting membrane proteins (2), each involving the mixing of a known amount of protein with a known amount of lipids, followed by a procedure to allow for the faithful incorporation of the protein into the bilayer while preserving the protein structure. Since most membrane proteins do not spontaneously insert into lipid bilayers (3), this requires a gentle approach of exchanging the protein from the stable detergent solubilized state to a lipid bilayer solvated condition. A common method to do this is to mix a small amount of protein stabilized in a detergent micelle, with lipids that have been solubilized with a detergent with a high critical micelle concentration. From this mixed-micelle condition, the detergent is then slowly removed by dialysis, resulting in the formation of bilayers with the protein incorporated within, that then assemble into self-contained liposomes (4).

One of the benefits of the compartmentalized structure of liposomes is that it enables the establishment of gradients across the membrane, which allows interrogation of the protein’s function (5, 6). One limitation, is that dialysis methods typically generate small, unilamellar vesicles (7) that incorporate the protein according to the state in the mixed micelle condition. Since liposomes do not spontaneously exchange with one another on a reasonable time scale, this means that the protein state that is examined in the resultant proteoliposome population may not reflect equilibrium. This presents an issue for considering stoichiometry of the protein, particularly at ‘Poisson-dilutions’ where most of the liposomes are unoccupied or have a single protein species. However, this is the range of dilution that is most desirable when considering protein function, as the membrane transport behavior reflects a single-molecule and so unitary transport rates can be estimated (8). Still, when investigating the function of the native assembly, it is advantageous to capture the equilibrium state of the protein in the membrane into single vesicles for further analysis of function and dynamics.

One way of doing this is by taking the small unilamellar liposomes from the dialysis procedure and fusing them into larger membranes. It has been observed that repeated cycles of freezing and thawing can lead to the growth of large multilamellar, or multivesicular, membranes with increased trapped volume (9, 10). This occurs due to the formation of ice during freezing which break the membrane structure, allowing for annealing with other membranes. For *E. coli* polar lipids, phase contrast optical microscopy showed that the freeze-thaw process forms large multilamellar vesicles with outer diameters of 10 µm (11), much larger than a typical biological cell membrane. In these large membranes, it now becomes possible for protein species to encounter one another and sample the thermodynamic consequences of the system. From here, smaller liposomes can be formed by fractionating the membrane by extrusion through polycarbonate filters with defined pore sizes to produce reproducible distributions of liposomes. During this process, the large membrane is pushed through the pores, anywhere from 30 nm to 1 µm, forming a membrane bubble that breaks off into a vesicle that is approximate in diameter to the extrusion pore size (12). The probability of protein incorporation into that vesicle is a random process that depends on the density of protein in the overall membrane, and the number of vesicles in the resultant liposome population (8, 13, 14).

It is advantageous to understand how protein is incorporated into the liposome population, for the interpretation of functional and structural studies as described above. If the liposome size distribution is uniform and there is a single protein species of interest, then this can be calculated using the formula for the Poisson distribution (equation 1). However, realistically, extruded liposomes lead to non-uniform distributions except for the smallest pore sizes. Liposome size distributions can be directly measured by imaging liposome samples, using freeze-fracture or cryo-electron microscopy (EM) approaches (7). Alternatively, numerical distributions can be obtained by dynamic light scattering spectroscopy, which exhibits good agreement with EM studies (15). These studies have revealed that extrusion of freeze-thawed multilamellar membranes through polycarbonate filters leads to vesicle populations with mean diameters comparable to the pore size of the filter (16). However, there is significant polydispersity when using pore sizes > 200 nm, often yielding bimodal distributions. In particular, 400 nm extruded liposomes obtained from freeze-thawed *E. coli* polar lipids or 2:1 1-palmitoyl-2-oleoyl-sn-glycero-3-phosphoethanolamine (POPE):1-palmitoyl-2-oleoyl-sn-glycero-3-phospho-(1’-rac-glycerol) (POPG) membranes, are much smaller than expected, with a peak population near 25 nm and a long rightward tail of larger vesicles (8, 17). The heterogeneity in liposome sizes has an important impact on the reconstitution statistics of membrane proteins, which must be known for proper interpretation of experimental data, in particular for knowing the ‘Poisson-dilution’ range, and the expected oligomeric assemblies during equilibrium reactions.

In this study, we examine the effect of liposome heterogeneity on the statistics of membrane protein reconstitution into liposomes. First, we outline the theory behind calculating the liposome occupancy probability distribution for heterogeneous liposome populations. We focus on the 2:1 POPE:POPG lipid composition, a synthetic mimic of the *E. coli* membrane and a commonly used lipid composition in reconstitution studies. By comparing simulations with experimental data, we show that the heterogeneous profile of the 400 nm extruded population, agrees with previously published data on fractional occupancy by co-localization microscopy, and fractional volume measured by proteoliposome efflux assays. Next, we measure the size distributions of 2:1 POPE:POPG extruded with 30 nm and 100 nm pores, and demonstrate that the single-molecule photobleaching distributions of the CLC-ec1 Cl^-^/H^+^ antiporter labeled with Cy5 agrees with simulations based on these liposome distributions. Finally, we show how extrusion through 30 nm filters increases the ‘Poisson-dilution’ limit and expands the dynamic range of discriminating between expected monomer and dimer reconstitutions distributions. These results may be useful in future experiments examining CLC-ec1 dimerization equilibrium, or studies of other membrane protein oligomers in membranes.

## 2. METHODS

### 2.1 MATLAB simulations

MATLAB was used to simulate the random process of multi-species subunit capture into a heterogeneous liposome population, as described previously (14, 18). For each liposome sub-population with radius *r*, a matrix was created with size *N*_*liposomes*_*(r)*, and each *N*_*protein*_*(r)* sub-species was randomly assigned to the liposome population. Unless otherwise noted, the following parameters were used: area per lipid, *A*_*lipid*_ = 0.6 nm^2^ (19); probability of subunit labeling, *P*_*Cy5*_ = 0.7; probability of non-specific subunit labeling, *P*_*ns*_ = 0.1; mole-fraction recovery, *yield* = 0.5; liposome size distribution, *P*_*radius*_*(r)*; liposomes accessibility factor, *bias* = 4 (excludes 2.5–22.5 nm bins from *P*_*radius*_*(r)*), to model the exclusion of ∼(5 nm × 10 nm) CLC-ec1 dimers from smaller liposomes. MATLAB files for the simulation program, and the analytical solution of the heterogeneous Poisson distribution are included as supplementary information.

### 2.2 Cryo-electron microscopy imaging of liposomes

A 2:1 mixture of 1-palmitoyl-2-oleoyl-sn-glycero-3-phosphoethanolamine: 1-palmitoyl-2-oleoyl-sn-glycero-3-phospho-(1’-rac-glycerol) (POPE:POPG (Avanti Polar Lipids Inc., Alabaster AL)) in chloroform was dried under a N_2_ gas line whilst the vial was slowly spun by hand to ensure an even distribution. The sample was then resuspended in 1 mL of pentane before being dried again. Dialysis buffer (300 mM KCl, 20 mM citric acid, pH 4.5 using NaOH) was added to the lipids to a final concentration of 20 mg/mL. Next, 35 mM of 3-((3-cholamidopropyl) dimethylammonio)-1-propanesulfonate (CHAPS (Anatrace, Maumee OH)) detergent was added and the solution was sonicated in a cylindrical bath sonicator for 30–60 minutes, until the cloudy solution had become transparent. The sample was dialyzed in Slide-A-Lyzer cassettes (Thermo Scientific, Waltham MA), 10 kDa MWCO, for 2 days at 4°C in dialysis buffer which was changed every 10–12 hours. Once dialysis was complete, the empty liposomes were freeze-thawed seven times by incubating in a dry ice/ethanol bath for 5 min, followed by a room temperature water bath for 15 min, before being stored at room temperature until they were required for imaging. Note, thawing occurs above the phase transition temperature, previously reported to be ∼19 °C (11). This procedure results in large multilamellar vesicles that are turbid and settle at the bottom of the tube (Figure 4A). On the day of imaging, liposomes were extruded 21-times through filters with pore sizes of 30, 100 and 400 nm then applied to glow-discharged Lacey carbon support films (Electron Microscopy Sciences, Hatfield PA) in 3 µL drops, blotted, and plunged into liquid ethane using a Vitrobot system (FEI). Grids were imaged at 300 kV in a JEOL 3200 fs microscope with K2 Summit direct electron detector camera (Gatan). A magnification of 30,000 was used for the 30 nm and 100 nm samples, and a magnification of 15,000 used for the 400 nm sample. For analysis, the polygon selection tool in ImageJ (20) was used to determine the radius of each liposome in nanometers in order to produce radius distributions. Note, all liposomes were analyzed include those lying on the carbon support. For each extrusion pore size, two distributions were obtained: an outer radius distribution and a total radius distribution. The outer radius assumed all liposomes were unilamellar and so only recorded the radius of the outmost liposome in multilamellar liposomes. On the other hand, the total radius was the sum of all radii in multilamellar liposomes meaning the distribution was stiffed slightly to the right with some liposomes having much larger radii.

### 2.3 Protein purification and reconstitution

The methods of protein purification were carried out as described previously (14, 17). Purified C85A/H234C CLC-ec1 with a C-terminal hexahistidine tag in 150 mM NaCl, 20 mM 3-(*N*-morpholino)propanesulfonic acid (MOPS), pH 7.0 and 5 mM n-Decyl-β-D-Maltopyranoside, DM, (Anatrace, Maumee OH) was labeled with Cy5-malemide (Lumiprobe, Hunt Valley MD) for 10 minutes followed by quenching with 100 mM cysteine. Free dye was removed by binding the protein to a cobalt affinity column, with successive washes. The labeled protein was eluted with 400 mM imidazole, and then the eluate was run on a ∼2.5 mL Sephadex G-50 (GE Lifesciences) size exclusion column to separate the imidazole and allow for quantification of the protein labeling yield by measuring the absorbances at 280 nm and 655 nm. The average labeling yield was *P*_*Cy5*_ = 0.65 ± 0.01 (mean ± std, n = 3). The fluorescently labeled protein in DM micelles was added to 20 mg/mL 2:1 POPE:POPG solubilized in 300 mM KCl, 20 mM citrate, pH 4.5 adjusted with NaOH and 35 mM CHAPS. The samples were dialyzed in 10 kDa MWCO cassettes, against a 2000-fold excess volume of the same buffer without the detergent, in the dark, at 4°C, with 3–4 buffer changes overall, with changes every 8–12 hours. The last round of dialysis was carried out at room temperature.

**Table 1:** 2:1 POPE:POPG extruded vesicle distributions – outer radii.

| *r* (nm) | *SA* (nm<sup>2</sup>) | *30 nm* |  | *100 nm* |  |
| --- | --- | --- | --- | --- | --- |
|  |  | *P<sub>radius</sub>* | *F<sub>SA</sub>* | *P<sub>radius</sub>* | *F<sub>SA</sub>* |
| 0-5 | 7.85E+01 | 0 | 0 | 0 | 0 |
| 5-10 | 7.07E+02 | 0.001 | 0.000 | 0.047 | 0.001 |
| 10-15 | 1.96E+03 | 0.037 | 0.005 | 0.109 | 0.008 |
| 15-20 | 3.85E+03 | 0.137 | 0.039 | 0.047 | 0.007 |
| 20-25 | 6.36E+03 | 0.186 | 0.087 | 0.050 | 0.012 |
| 25-30 | 9.50E+03 | 0.200 | 0.140 | 0.103 | 0.036 |
| 30-35 | 1.33E+04 | 0.147 | 0.142 | 0.080 | 0.039 |
| 35-40 | 1.77E+04 | 0.097 | 0.126 | 0.059 | 0.038 |
| 40-45 | 2.27E+04 | 0.075 | 0.126 | 0.088 | 0.074 |
| 45-50 | 2.84E+04 | 0.044 | 0.092 | 0.079 | 0.083 |
| 50-55 | 3.46E+04 | 0.039 | 0.099 | 0.076 | 0.097 |
| 55-60 | 4.15E+04 | 0.014 | 0.044 | 0.053 | 0.081 |
| 60-65 | 4.91E+04 | 0.012 | 0.042 | 0.070 | 0.127 |
| 65-70 | 5.73E+04 | 0.008 | 0.033 | 0.068 | 0.142 |
| 70-75 | 6.61E+04 | 0.002 | 0.011 | 0.024 | 0.057 |
| 75-80 | 7.55E+04 | 0.001 | 0.004 | 0.015 | 0.041 |
| 80-85 | 8.55E+04 | 0 | 0 | 0.009 | 0.028 |
| 85-90 | 9.62E+04 | 0.001 | 0.006 | 0.006 | 0.021 |
| 90-95 | 1.08E+05 | 0.001 | 0.006 | 0.006 | 0.023 |
| 95-100 | 1.19E+05 | 0 | 0 | 0 | 0 |
| 100-105 | 1.32E+05 | 0 | 0 | 0.003 | 0.014 |
| 105-110 | 1.45E+05 | 0 | 0 | 0.006 | 0.031 |
| 110-115 | 1.59E+05 | 0 | 0 | 0 | 0 |
| 115-120 | 1.73E+05 | 0 | 0 | 0.003 | 0.019 |
| 120-125 | 1.89E+05 | 0 | 0 | 0 | 0 |
| 125-130 | 2.04E+05 | 0 | 0 | 0.003 | 0.022 |
| 130-135 | 2.21E+05 | 0 | 0 | 0 | 0 |
| 135-140 | 2.38E+05 | 0 | 0 | 0 | 0 |
| 140-145 | 2.55E+05 | 0 | 0 | 0 | 0 |
| 145-150 | 2.73E+05 | 0 | 0 | 0 | 0 |
| 150-155 | 2.92E+05 | 0 | 0 | 0 | 0 |
| 155-160 | 3.12E+05 | 0 | 0 | 0 | 0 |
| 160-165 | 3.32E+05 | 0 | 0 | 0 | 0 |
| 165-170 | 3.53E+05 | 0 | 0 | 0 | 0 |
| 170-175 | 3.74E+05 | 0 | 0 | 0 | 0 |
| 175-180 | 3.96E+05 | 0 | 0 | 0 | 0 |
| 180-185 | 4.19E+05 | 0 | 0 | 0 | 0 |
| 185-190 | 4.42E+05 | 0 | 0 | 0 | 0 |
| 190-195 | 4.66E+05 | 0 | 0 | 0 | 0 |
| 195-200 | 4.90E+05 | 0 | 0 | 0 | 0 |
| 200-205 | 5.15E+05 | 0 | 0 | 0 | 0 |
| 205-210 | 5.41E+05 | 0 | 0 | 0 | 0 |
| 210-215 | 5.67E+05 | 0 | 0 | 0 | 0 |
| 215-220 | 5.94E+05 | 0 | 0 | 0 | 0 |
| 220-225 | 6.22E+05 | 0 | 0 | 0 | 0 |
| 225-230 | 6.50E+05 | 0 | 0 | 0 | 0 |
| 230-235 | 6.79E+05 | 0 | 0 | 0 | 0 |
| 235-240 | 7.09E+05 | 0 | 0 | 0 | 0 |
| 240-245 | 7.39E+05 | 0 | 0 | 0 | 0 |
| 245-250 | 7.70E+05 | 0 | 0 | 0 | 0 |
| 250-255 | 8.01E+05 | 0 | 0 | 0 | 0 |
| 255-260 | 8.33E+05 | 0 | 0 | 0 | 0 |
| 260-265 | 8.66E+05 | 0 | 0 | 0 | 0 |
| 265-270 | 8.99E+05 | 0 | 0 | 0 | 0 |
| 270-275 | 9.33E+05 | 0 | 0 | 0 | 0 |
| 275-280 | 9.68E+05 | 0 | 0 | 0 | 0 |
| 280-285 | 1.00E+06 | 0 | 0 | 0 | 0 |
| 285-290 | 1.04E+06 | 0 | 0 | 0 | 0 |
| 290-295 | 1.08E+06 | 0 | 0 | 0 | 0 |
| 295-300 | 1.11E+06 | 0 | 0 | 0 | 0 |

**Table 2:**
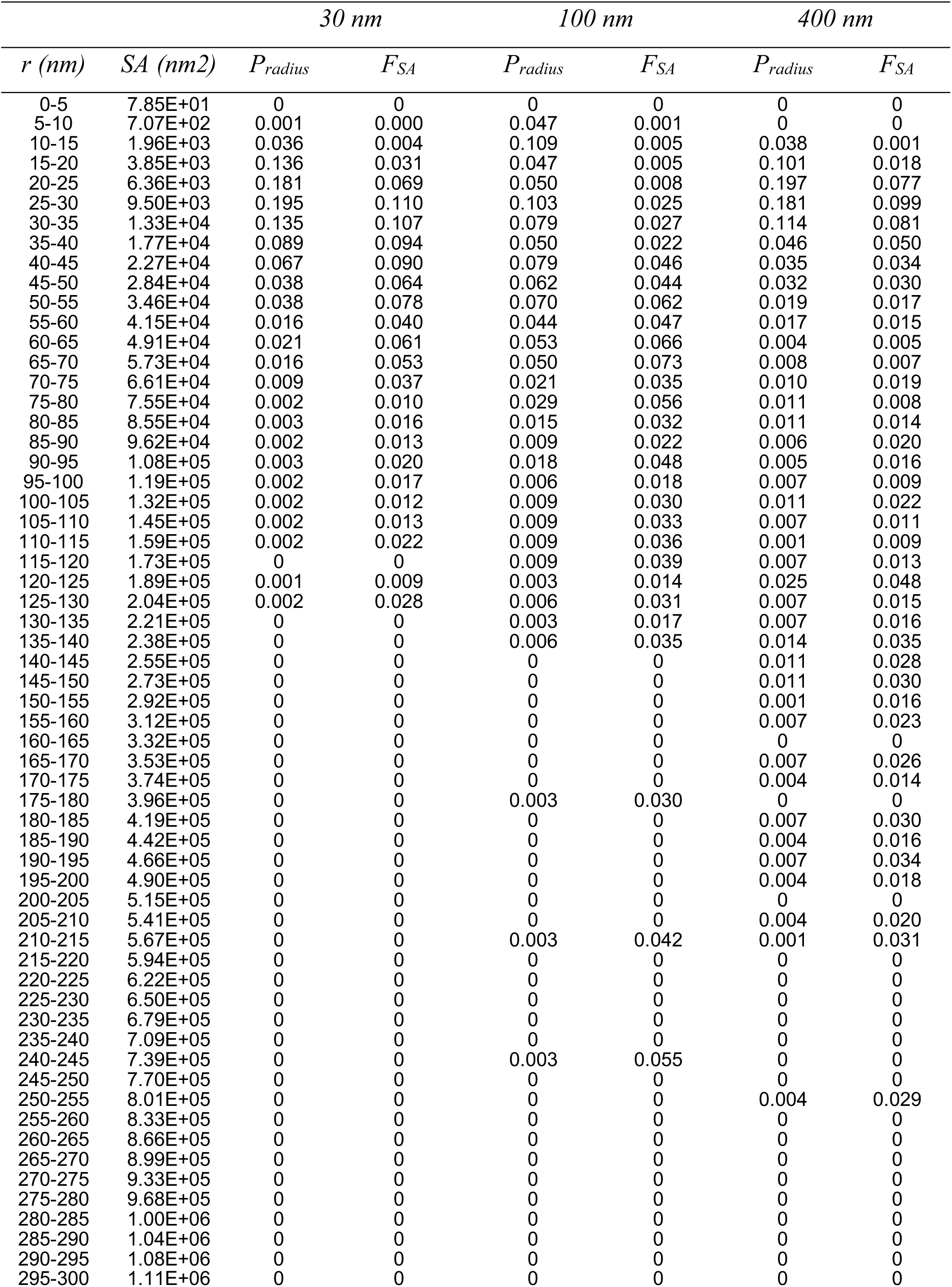
POPE:POPG extruded vesicle distributions ℃ total radii.

Calculating the reconstitution density is carried out as follows. CLC-ec1 C85A/H234C including the hexahistidine tag has a molecular weight of 51,997 g/mole, *MW*_*POPE*_ = 717.996 g/mole and *MW*_*POPG*_ = 770.989 g/mole. Therefore, a proteoliposome sample reconstituted at 1 µg/mg in 2:1 POPE/POPG lipids corresponds to a mole fraction of:

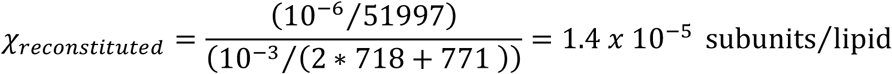

Assuming a 50% mole fraction recovery as was previously observed (14), then our observed mole fraction is *χ*_*observed*_ = 0.7 *x* 10^−5^ subunits/lipid.

### 2.4 Single-molecule TIRF microscopy of fluorescently labelled proteoliposomes

#### Co-localization microscopy

Studies were carried out as previously described (14). Briefly, the primary amine on POPE was conjugated to Alexa Fluor 488 Carboxylic Acid, 2,3,5,6-Tetrafluorophenyl (TFP) ester (Thermo Fisher Scientific) during the POPE/POPG/CHAPS stage, at room temperature and pH 8.0. After 4 hours of incubation, excess free dye was quenched with Tris, and then the lipids were used in the reconstitution procedure as usual, with free dye removed during dialysis. After freeze-thaw and extrusion, liposomes were applied to the slide, and first imaged with the longer wavelength 637 nm laser to detect the Cy5 labeled protein. Next, we imaged the liposomes in the same field using the 488 nm laser. Multicolor tetraSpeck 0.1 µm beads (Thermo Fisher Scientific) or co-labeled liposomes were loaded onto a separate lane for imaging in both channels, in order to create an independent mapping file to determine co-localization using a custom-built MATLAB software (21).

#### Photobleaching analysis

Photobleaching analysis was carried out as described previously (14). After completion of dialysis, multilamellar vesicles were obtained by seven cycles of freezing (dry-ice/ethanol bath for 5 min or −80 °C, 10 minutes) and thawing (room temperature water bath, 10–15 minutes) of the proteoliposome samples. Sodium azide (0.02% w/v) was added to each sample which was then incubated at room temperature for 3–4 days before imaging. Immediately prior to imaging, proteoliposomes were sequentially extruded using 0.4, 0.1, and 0.03 micron nucleopore filters (GE Lifesciences, Maidstone UK) using a LiposoFast Basic extruder (Avestin Inc., Ottawa, Canada). Imaging of fluorescent spots and analysis of steps taken before complete photobleaching was performed as described before (14).

## 3. Theory

When starting from the large multilamellar membrane state, the act of capturing protein into vesicles as they are extruded can be modeled as a spatial Poisson process. In reality, this is a nonhomogeneous Poisson process because of the diversity of the types of protein species that may be in the membrane, as well as heterogeneity in the surface area of liposomes that form the compartments. Here, we focus on outlining the impact of liposome polydispersity on the resultant membrane protein occupancy distribution.

### 3.1 Liposome occupancy distribution with homogenous compartments

First consider the Poisson distribution for a homogenous system with a single protein species and one size of vesicle with radius *r*. The distribution is calculated as,

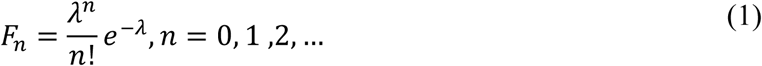

where *n* is the protein occupancy value, and the *λ* is the Poisson parameter,

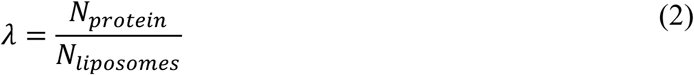

which is equal to the distribution mean and variance. In reconstitution experiments, an experimentalist explicitly sets the number of protein particles, *N*_*protein*_, and number of lipids, *N*_*lipids*_, in a sample for a defined density, or mole fraction of protein per lipid, *χ*,

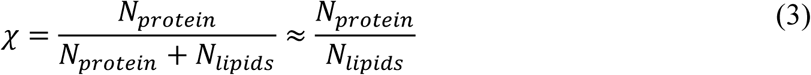

For many lipids, the surface area of a lipid in the bilayer, *A*_*lipid*_, has been experimentally determined (22), and so the total bilayer surface area, *SA*_*total*_, can be calculated as:

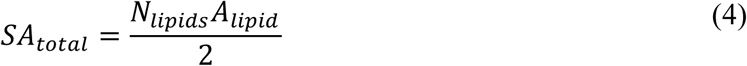

With this, *N*_*liposomes*_ can be determined,

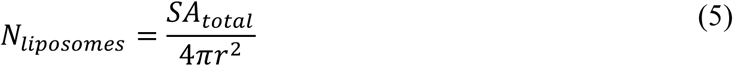

and substituting equations (3–5) into (2) gives,

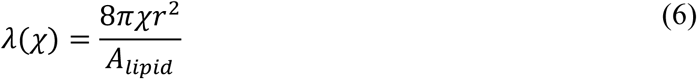

### 3.2 Liposome occupancy distribution with heterogeneous compartment

For a heterogeneous population of liposomes with a single protein species, the reconstitution distribution can still be solved for analytically as long as the probability distribution of liposomes sizes, *P*_*radius*_*(r)*, is known. Here, we consider that each sub-population of liposomes with radius *r*, reduces to an independent homogeneous Poisson process with a different Poisson parameter that depends on the radius of the sub-population:

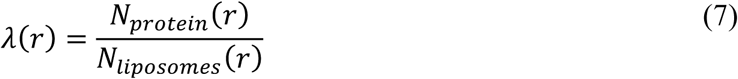

Note, the number of liposomes depends on the radius for the probability of occurring in the population (Figure 1A). The number of protein particles also depends on the radius as the protein is allocated proportional to the fraction of the total membrane area in that sub-population (Figure 1B).

**Figure 1.**
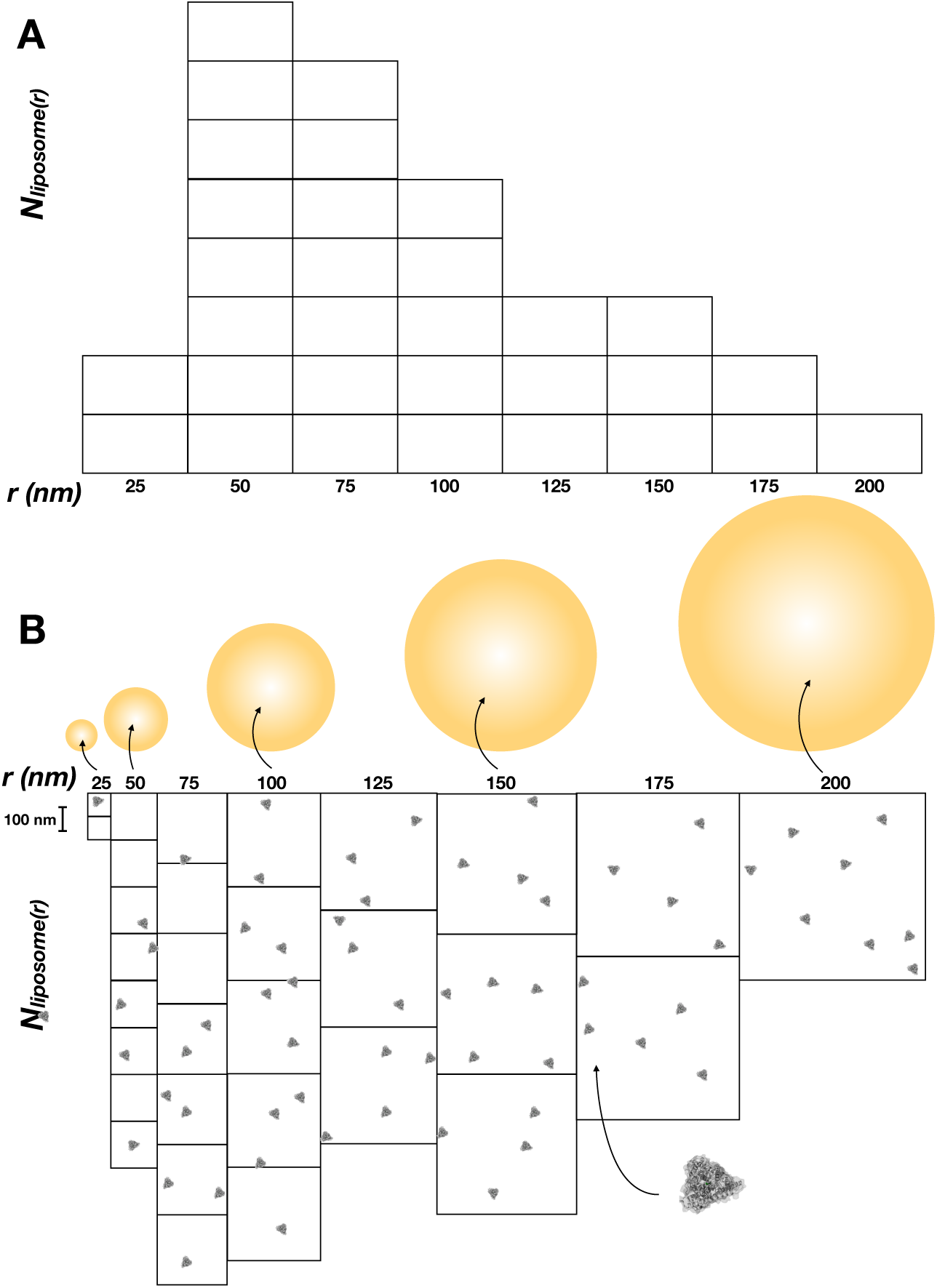
The effect of liposome heterogeneity on membrane protein capture statistics in liposomes. (A) A hypothetical heterogeneous size distribution of liposomes. Normalization of this distribution by the total number of liposomes gives *P*_*radius*_*(r)* (B) The same distribution drawn as a scaled representation of the liposome surface area, *SA* = 4*πr*^2^. Overlay of randomly distributed CLC-ec1 monomers (enlarged to be visible) over the total membrane area, corresponding to a density of *χ*_*reconstituted*_ ∼ 5 *x* 10^−6^ subunits/lipid (i.e. ∼0.5 µg/mg for CLC-ec1 subunit). This cartoon illustrates how larger vesicles have an increased probability of being being occupied due to their increased allocation of membrane area. In contrast, smaller vesicles have a higher likelihood to remain unoccupied at similar densities.

The occupancy distribution for each sub-population is therefore,

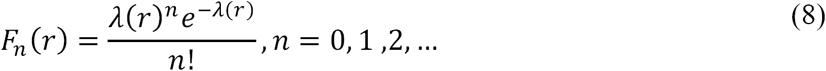

and the resultant occupancy distribution across all liposomes is:

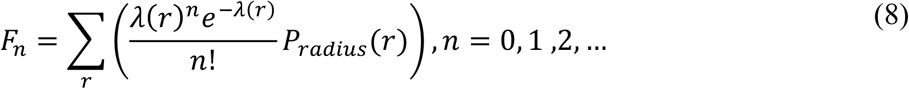

The radius dependent Poisson parameter is equivalent to the homogenous form derived in equation (6). This is shown as follows:

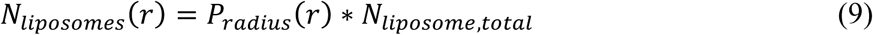

Where *N*_*liposome,total*_ is the total number of liposomes in the population. To calculate this, we need to again consider the total bilayer surface area set in the experiments:

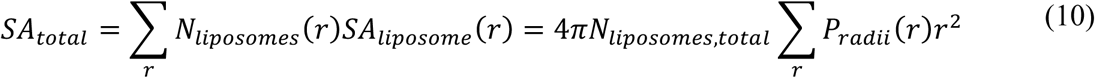

Equating this to equation (4) allows us to solve the total number of liposomes in the sample in a heterogeneous population:

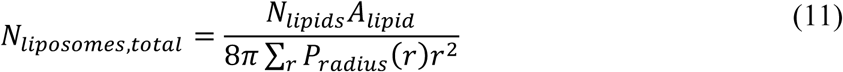

Thus, the number of liposomes in each sub-population of radius r is calculated as,

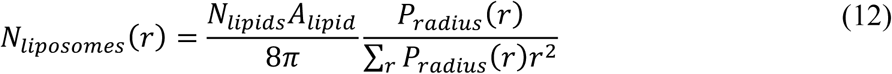

The number of protein particles that are available to partition into each sub-population of liposomes, is proportional to the fraction of membrane surface area occupied by *N*_*liposomes*_*(r)*.

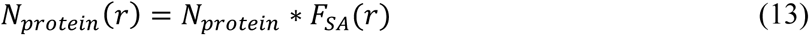

Where the fractional surface area is equal to:

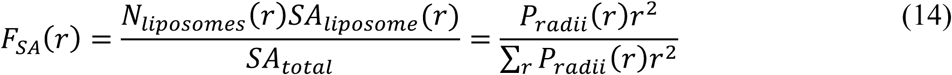

Therefore,

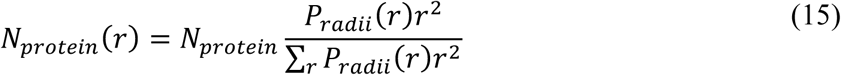

Substituting (15) and (12) into (7) reduces to equation (6), but with variable *r*:

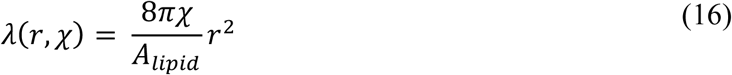

### 3.3 Numerical fraction of unoccupied vesicles

The fraction of unoccupied vesicles, *F*_*0,num*_, can simply be calculated for n = 0:

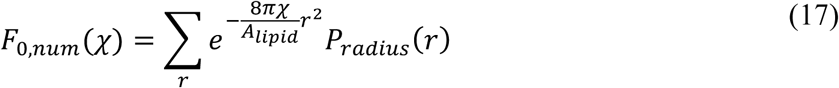

For the homogenous distribution, this reduces to a single exponential decay:

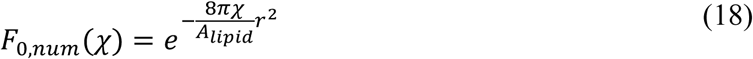

### 3.4 Fractional volume of unoccupied vesicles

The fraction of unoccupied vesicle volume can also be calculated, *F*_*0,vol*_. First, consider the volume trapped inside of the sub-population of liposomes with radius *r, V*_*liposomes*_*(r)*:

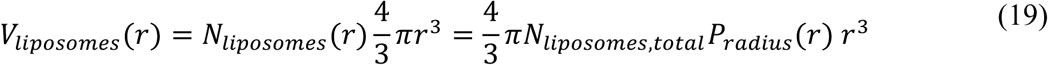

And the total volume encapsulated in all of the liposomes,

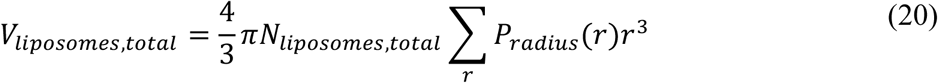

Therefore, the fractional volume is:

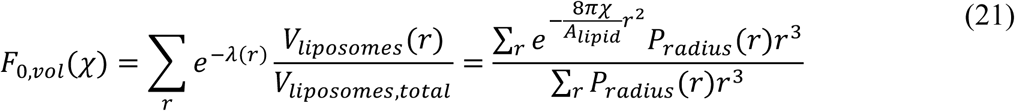

### 3.5 Simulations of membrane protein reconstitution

While the analytical solutions above allow for simple calculations of single species incorporated into heterogeneous liposome populations, the calculations become more complicated once multiple protein species are introduced. This requires enumeration of the distribution of different species, and treatment of each as an independent Poisson process into the same liposome population. The overall occupancy then depends on many joint probabilities of co-occupancy of various protein species, of which there are many combinations, making the calculation daunting. A simpler solution is offered by statistical simulation in MATLAB that carries out the random assignment of protein species into matrices representing different liposome sub-populations. Figure 1 shows a schematic of the simulation, which highlights the impact of the differential surface area in a heterogeneous distribution. The details of this method have been outlined previously and we refer the reader to these papers for futher details on the simulation(14, 17). The key parameters in the simulation are all experimentally determined, including the area per lipid, *A*_*lipid*_ = 0.6 nm^2^ (19); probability of subunit labeling, *P*_*fluor*_; probability of non-specific subunit labeling, *P*_*ns*_ measured in a cysteine-inaccessible construct; mole-fraction recovery, *yield*; liposome size distribution, *P*_*radius*_*(r)*; and liposomes accessibility factor, *bias* = 4 (excludes 2.5–22.5 nm bins from *P*_*radius*_*(r)*), to model the exclusion of ∼(5 nm × 10 nm) CLC-ec1 dimers from smaller liposomes. MATLAB files for the simulation program, and the analytical solution of the heterogeneous Poisson distribution are included as supplementary information.

## 4. Results

### 4.1 The effect of liposome heterogeneity on occupancy statistics

For single-component membranes, it is generally assumed that the liposome population obtained from freeze-thaw/extrusion generates a uniform population with diameters matching the size of the extrusion pore (Figure 2A). Indeed, cryo-electron microscopy studies of 1-palmitoyl-2-oleoyl-glycero-3-phosphocholine (POPC) liposomes extruded through 50 nm pores show a distribution with a mean of 27 nm (23). In addition, egg PC vesicles extruded through 400 nm pores is reported to be “relatively homogeneous” with a distribution centered at 243 ± 91 nm (Figure 2B), as measured by freeze-fracture electron microscopy (12). On the other hand, lipid mixtures of *E. coli* polar lipid (EPL) extracts (8), or the synthetic mimic 2:1 POPE:POPG (17) yield a heterogenous distribution with a peak near 25 nm and a rightward tail towards larger liposomes (Figure 2C). The immediate consequence of the smaller liposomes means that there are ∼100-fold more liposomes in the 400 nm 2:1 POPE:POPG population than expected (Figure 2D). To examine how this affects the reconstitution statistics, we calculated the liposome occupancy distribution for the Poisson process of protein incorporation into liposomes, based on three examples of 400 nm extruded liposome size distributions: the idealized homogenous population, and experimental distributions of egg PC or 2:1 POPE:POPG. The fraction of occupied vesicles is similar for the ideal and egg PC conditions (Figure 2E) even though the widths of the two distributions are significantly different. Yet, the fraction of liposomes that are occupied does not change because the number of liposomes is comparable in the two conditions (Figure 2D). In contrast, the 2:1 POPE:POPG curve is shifted to the right, requiring much higher protein densities to saturate the vesicle population, a consequence of the increase in *N*_*liposomes*_. On the other hand, the fractional volume of empty vesicles is similar for the three distributions (Figure 2F). The fractional volume is often measured during functional measurements of ion efflux from vesicles, and has been used to inform on membrane protein reconstitution statistics and protein stoichiometry (8, 13, 24). Here, we see that the fractional volume is less sensitive to the changes in the liposome size distribution. Examining the protein occupancy statistics demonstrates how heterogeneity in the 2:1 POPE:POPG distribution leads to a spreading of single and double occupancy states over a wider range of densities (Figure 2G,H), but the increased number of liposomes reduces the probability of saturation with more than three protein subunits (Figure 2I). Note that for all cases, we have calculated the analytical solution in parallel to simulating the reconstitution process and there is complete agreement between the two methods.

**Figure 2.**
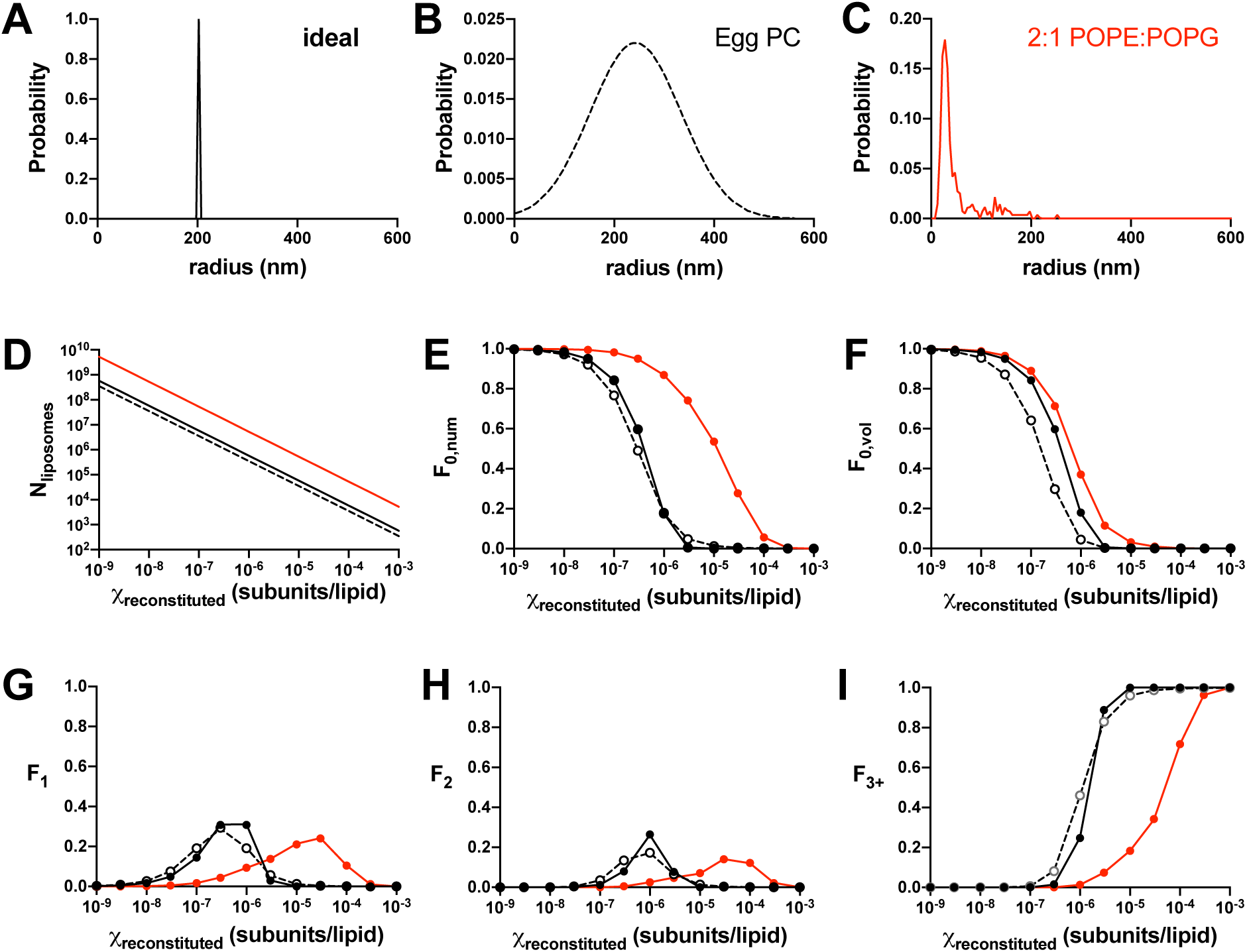
Reconstitution statistics into 400 nm extruded liposome populations. Calculations of the Poissonian reconstitution statistics of independent protein subunits into liposome populations with varying heterogeneity. (A) Model of ideally extruded, homogenous population with radius = 200 nm (black, solid). (B) Egg PC liposomes (12) following a gaussian distribution with µ = 243 nm and σ = 91 nm (black, dashed). (C) 2:1 POPE:POPG liposomes (Chadda *et al.*, 2018), demonstrating smaller than expected sizes and a rightward tail (red). (D) *N*_*liposomes*_ as a function of the reconstitution density (constant *N*_*protein*_ = 10^6^). (E) Fraction of unoccupied vesicles, *F*_*0,num*_. (F) Fractional volume of unoccupied vesicles, *F*_*0,vol*_, corresponding to liposomal efflux studies. Fraction of vesicles containing (G) one, (H) two, or (I) more than three subunits. The calculations were performed in two ways: analytical solution – circles, simulation – lines.

### 4.2 Testing the 400 nm 2:1 POPE:POPG liposome size distribution

The previous findings that EPL and 2:1 POPE:POPG lipid bilayers extruded through 400 nm pores yield significant number of liposomes with radius ∼ 25 nm is unexpected. To test whether this population agrees with experimental observations, we compared co-localization microscopy data of ‘WT’ CLC-ec1-Cy5 (i.e. C85A/H234C background) with AlexaFluor-488 labeled 400 nm extruded 2:1 POPE:POPG vesicles to the expected fraction of Cy5 unoccupied vesicles, \**F*_*0,num*_, simulated for the 400 nm extruded egg PC or 2:1 POPE:POPG liposome distributions (Figure 3A). Note, \**F*_*0,num*_, differs *F*_*0,num*_ because it includes liposomes that are occupied by unlabeled protein. The experimental co-localization data agrees with the 2:1 POPE:POPG distribution, with 50% of the vesicles occupied at *χ*_0.5_ ∼ 9 × 10^-5^ subunits/lipid while the egg PC distribution predicts a higher probability of filling at lower densities, with *χ*_0.5_ ∼ 2 × 10^-6^ subunits/lipid. Next, we compared the occupancy data from functional measurements of liposomal efflux of the CLC-ec1 Cl^-^/H^+^ antiporter (8) and Fluc F^-^ channel reported in the literature (13). In these “ion-dump” assays, liposomes are loaded with a high concentration of the transport ion, and then exchanged into a low ion concentration. Since the movement of the ion across the membrane is electrogenic for either protein, efflux is initiated by addition of valinomycin, which sets the membrane potential to zero and allows the ion to flow down its concentration gradient. Vesicles that do not contain active protein are still ion-loaded, and addition of a detergent that disrupts all membranes releases the remaining amount of trapped ion. From this, a fractional volume of empty vesicles, *F*_*0,vol*_, can be experimentally measured. We took three studies where *F*_*0,vol*_ was reported as a titration of protein density and extrapolated the data: WT CLC-ec1, E148A/Y445S CLC-ec1 and the EC2 Fluc F^-^ channel (8, 13). The data were compared to simulations of protein reconstitution into 400 nm extruded vesicles from egg PC or 2:1 POPE:POPG membranes (Figure 3B). Each set of experimental data overlays with one another, independent of the protein construct. Furthermore, the data agrees with the 2:1 POPE:POPG distribution, with *χ*_0.5_ ∼ 5 × 10^-6^ subunits/lipid compared to *χ*_0.5_ ∼ 7 × 10^-7^ subunits/lipid expected from an egg PC distribution. Altogether, both methods of measuring liposome occupancy agree with the presence of smaller liposomes indicated by the 2:1 POPE:POPG distribution, and not idealized or egg PC distributions.

**Figure 3.**
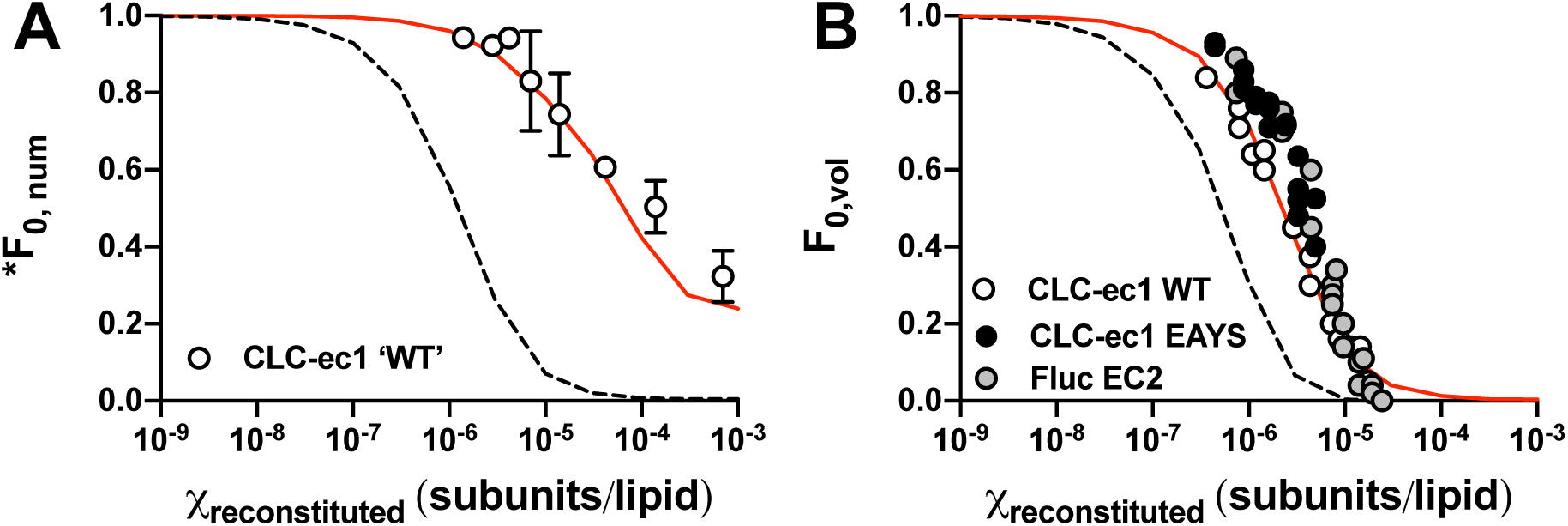
Experimental measurements of vesicle occupancy. Simulations of unoccupied vesicle fractions from 400 nm extruded 2:1 POPE:POPG liposomes (red) or 400 nm extruded egg PC liposomes (dashed). (A) Calculated fraction of vesicles that do not contain fluorescent protein, \**F*_*0,num*_. Parameters used: *P*_*fluor*_ = 0.7 & *P*_*ns*_ = 0.1, *yield* = 0.5, *K*_*D*_ = 2 × 10^-8^ subunits/lipids, and liposomes with *r* < 25 nm inaccessible to dimers. Co-localization microscopy experiments of ‘WT’, i.e. C85A/H234C CLC-ec1-Cy5 reconstituted in 2:1 POPE:POPG vesicles doped with POPE-AF488. Data shown as mean ± s.e.m, n = 3 ((14) and new data). (B) Calculated fractional volume of inactive vesicles, *F*_*0,vol*_, measured by the ion ‘dump’ assay, using the following parameters: *P*_*fluor*_ = 1 & *P*_*ns*_ = 0, *yield* = 0.5, *K*_*D*_ = 2 × 10^-8^ subunits/lipids, and liposomes with *r* < 25 nm inaccessible to dimers. Extrapolated data for WT CLC-ec1 (8), CLC-ec1 E148A/Y445S and Fluc EC2 (13) shown as symbols.

**Figure 4.**
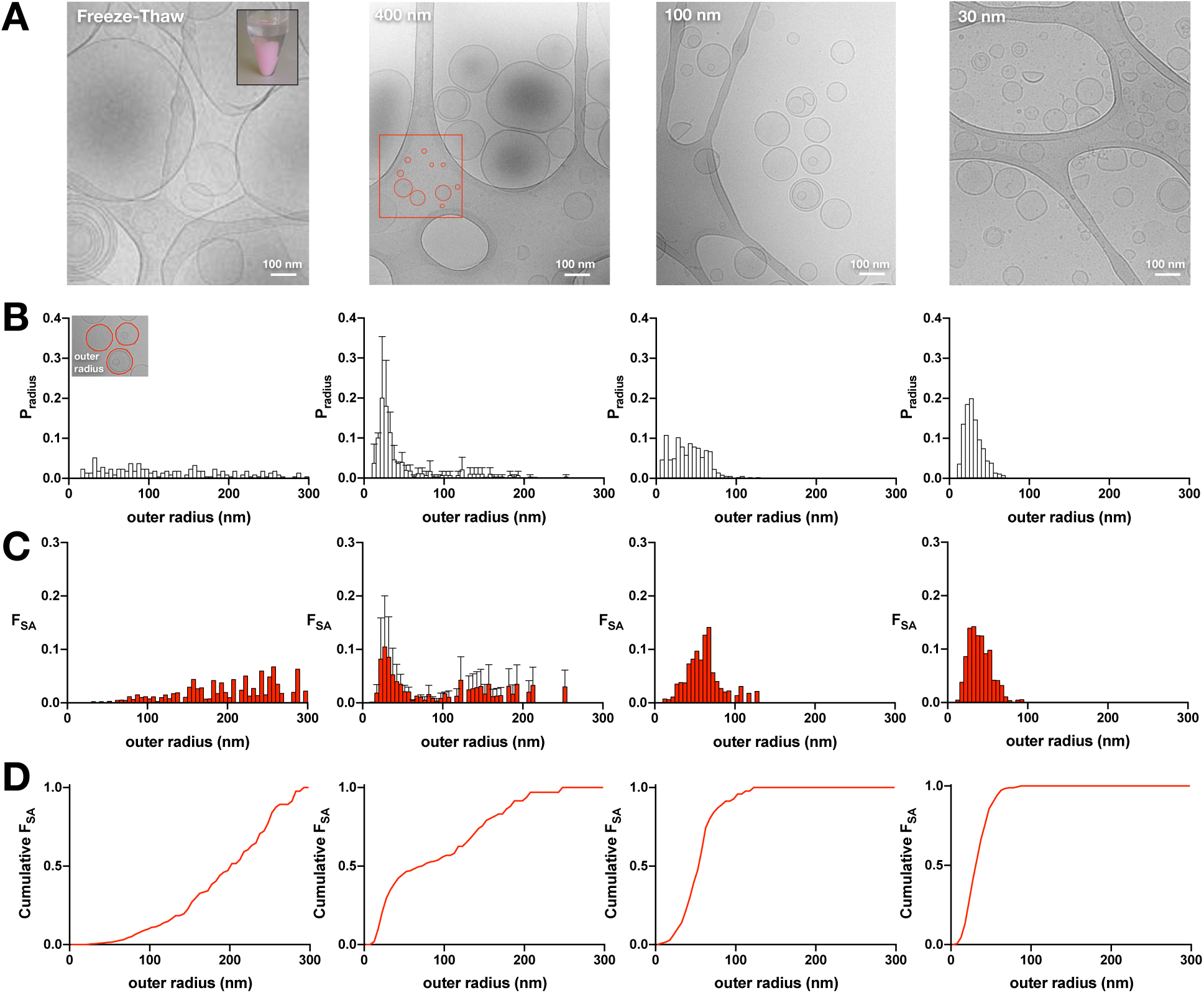
Size distributions of the outer radii of extruded liposomes composed of 2:1 POPE:POPG lipids. (A) Cryo-electron microscopy images of 2:1 POPE:POPG liposomes after seven cycles of freeze-thaw, or extrusion through 400 nm, 100 nm, or 30 nm filters. The inset in the freeze-thaw image shows the resultant large, multilamellar vesicles that are visible by eye and settle to the bottom of the tube (pink color is due to the addition of a small amount of POPE-RhodamineB). The red box in the 400 nm image highlights smaller vesicles identified on the carbon grid. (B) Probability distributions of the liposome outer radii, *P*_*radius*_, for the corresponding samples. (C) Probability distributions of the fractional surface area, *F*_*SA*_, for the corresponding samples. (D) Cumulative sum of *F*_*SA*_. The 30 nm and 100 nm distributions are presented in Table 1 while the 400 nm distribution was presented previously (17).

### 4.3 Liposome size distributions as a function of extrusion pore size

Considering the 400 nm 2:1 POPE:POPG liposomes are much smaller than expected, we went on to examine what happens when these membranes are extruded through even smaller pores. Figure 4A shows cryo-electron microscopy images of liposomes after freeze/thaw and following extrusion by 400 nm, 100 nm and 30 nm pores. The probability distributions of the outer radii of each vesicle (Figure 4B) and the fractional surface area (Figure 4C) are reported for each population. The mean ± standard deviation of outer radii of the freeze-thawed samples is 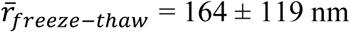 (and 48% as multilamellar vesicles). Note that the freeze-thawed distribution is not expected to be representative of the total population, as the larger vesicles that comprise the majority of the population, ∼ 10 µm in diameter (11) would not be captured in the thin layer of vitreous ice. The average values of the distributions decrease with extrusion, with 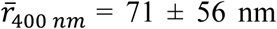 (13% MLVs), 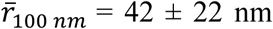 (17% MLVs) and 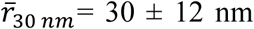 (10% MLVs), and they become more homogeneous. Note that the analysis included counting liposomes that resided on the carbon grid, outside of the vitreous ice for a complete representation of all vesicles in the population. The three extruded distributions of 2:1 POPE:POPG vesicles shows that the populations are not overwhelmingly different, and are all peaked around 25–30 nm. The major difference is the reduction of the larger liposome tail, resulting in increasingly homogenous populations of vesicles upon reducing the size of the extrusion pore. The fraction of MLVs in the extruded samples is small, 10–17% and does not change with pore size. To consider the increased amount of membrane in the MLVs, we also calculated the total radius distributions. Here, the area from all bilayers within a MLV are added together and counted as a single liposome with larger radius (Supplementary Figure 1). Since the fraction of multilamellar vesicles is small, this results in a nearly undetectable rightward shift of each distributions. The data for all of the liposome size distributions are reported in Tables 1 & 2.

### 4.4 Photobleaching distributions of protein occupancy

To examine how the reduction of liposome heterogeneity affects the reconstitution statistics, we carried out experiments of single-molecule photobleaching analysis of CLC-ec1-Cy5 in 2:1 POPE/POPG liposomes extruded through the three different pore sizes (Figure 5). At low densities, *χ*_*reconstituted*_= 10^-6^ & 10^-7^ subunits/lipid, there is no change in the photobleaching probability distributions across the different extrusions. This is likely because the protein is at densities below the ‘Poisson-dilution’ limit, where there is a near zero probability of a vesicle being occupied by more than one protein. However, this also shows that the probability distribution, and monomer-dimer populations, are independent of the extrusion method. On the other hand, extrusion of the highest density sample, *χ*_*reconstituted*_= 10^-5^ subunits/lipid, is dependent on the extrusion pore size. At this density, liposomes contain many protein particles, and so reducing the number of larger liposomes in the population allows the protein to distribute across more vesicles. A direct comparison of the simulations with this new experimental data (Supplementary Figure 2) shows the overall agreement and predictive power of the simulation, provided the liposome size distribution is experimentally measured.

**Figure 5.**
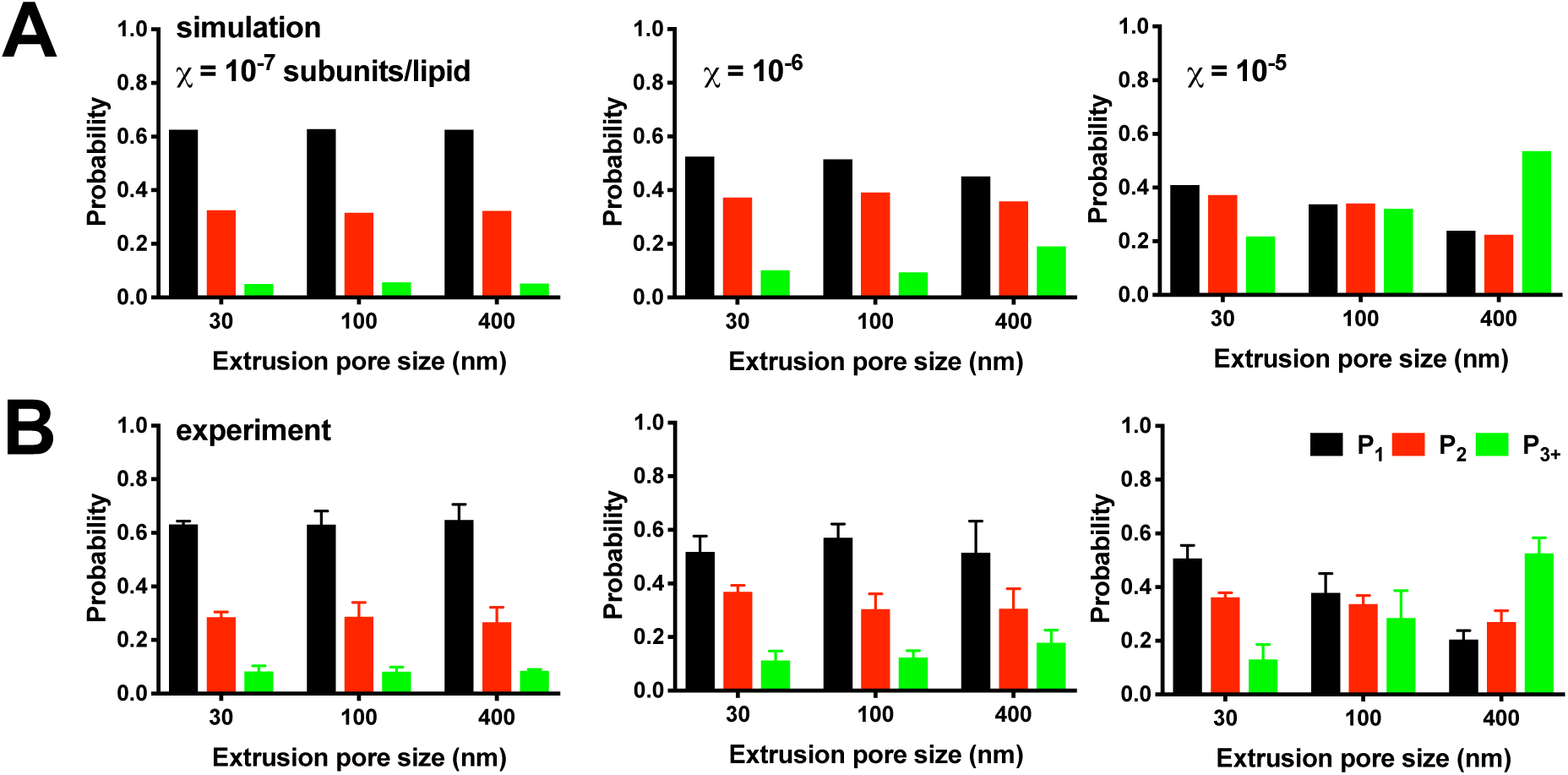
Photobleaching distributions as a function of liposome size. (A) Simulations of the expected (*P*_*1*_, *P*_*2*_, *P*_*3+*_) photobleaching distribution at *χ* = 10^-7^, 10^-6^ and 10^-5^ subunit/lipid densities, using the 2:1 POPE:POPG liposome size distributions presented in Figure 4. Simulation parameters include fluorophore labeling yields *P*_*fluor*_ = 0.7 & *P*_*ns*_ = 0.1, reconstitution yield = 1, monomer-dimer equilibrium *K*_*D*_ = 1 × 10^-8^ subunits/lipids, and liposomes with *r* < 25 nm inaccessible to dimers. (B) Experimental photobleaching distributions of ‘WT’ CLC-ec1-Cy5 in 2:1 POPE:POPG liposomes as a function of protein density and extrusion pore size. Data shown as mean ± s.e.m, n = 3–4.

### 4.5 30 nm extrusion expands the dynamic range of discriminating between monomer and dimer reconstitution statistics

Previously, we have shown that photobleaching analysis of subunit-capture statistics into liposomes can be used to inform on the populations of protein stoichiometry, from which information about equilibrium reactions in the membranes can be obtained (14). For instance, CLC-ec1 behaves in a monomer-dimer equilibrium that can be followed by changes in the photobleaching probability distribution as a function of the density. The power of this approach depends on the ability for protein species - monomers, dimers or other oligomeric forms - to be captured independently into different liposomes. Over-filling of liposomes at higher densities limits the ability to detect differences between protein populations limiting the dynamic range of the experiment. One possible solution is to use a smaller liposome population that will allow for the protein to distribute across more vesicles to allow for better resolution of the monomer vs. dimer statistics. Figure 6 shows the expected non-reactive monomer (Figure 6A) and non-reactive dimer (Figure 6B) occupancy statistics for the three distributions of 2:1 POPE:POPG liposomes. To calculate the fraction of dimer in the population, a least-squares method of estimation is used to fit the experimental data between the monomer and dimer benchmarks. Figure 6C shows the maximum signal for the least-squares analysis, *R*_*max*_, or distance between the monomer and dimer vectors, defined by the various combination of experimental values: *P*_*1*_, *P*_*2*_, *P*_*3+*_ and \**F*_*0,num*_. The 30 nm distribution leads to an increase in R_max_ at higher densities, increasing the power in estimating *F*_*Dimer*_ as a function of the mole fraction.

**Figure 6.**
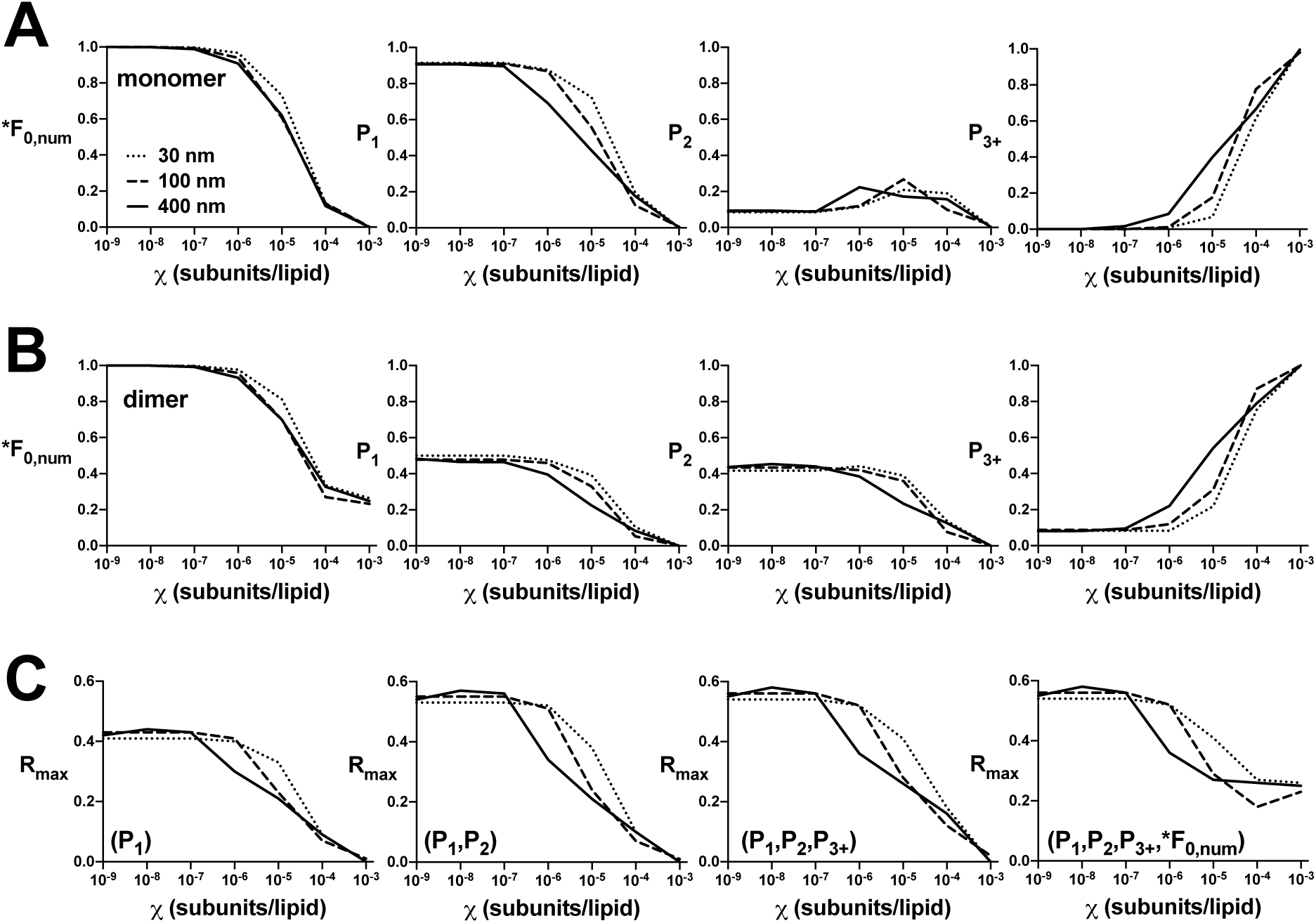
The dynamic range of single-molecule fluorescence microscopy measurements of subunits occupancy as a function of liposome size. Simulation of subunit capture of CLC-ec1 (fluorescent labeling yields: *P*_*fluor*_ = 0.7 & *P*_*ns*_ = 0.1) as a function of reconstituted protein density *χ* subunits/lipid, assuming 100% recovery (yield = 1). \**F*_*0,num*_ – fraction of unoccupied vesicles by co-localization microscopy. *P*_*1*_, *P*_*2*_, *P*_*3+*_ – single molecule photobleaching probabilities of fluorophore occupied vesicles. (A) non-reactive monomer (*K*_*D*_ = 10^100^ subunits/lipid). (B) non-reactive dimer (*K*_*D*_ = 10^-100^ subunits/lipid), liposomes r < 25 nm as inaccessible to protein. (C) Maximal least-squares range between expected monomer and dimer signals for fraction of dimer, *F*_*Dimer*_, estimation.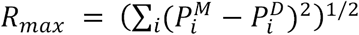 where *P*_*i*_ corresponds to *P*_*1*_, *P*_*2*_, *P*_*3+*_ of \**F*_*0,num*_.

## 5. Discussion

In this study, we examine how the liposome population, as a function of size and heterogeneity, affects the reconstitution statistics of membrane proteins into vesicles. Previously, we found that liposomes obtained by extruding freeze-thawed 2:1 POPE:POPG membranes through 400 nm filters generate a heterogeneous size distribution containing smaller liposomes with peak radii around 25 nm and a long tail of larger vesicles (17), in agreement with results reported for membranes derived from *E. coli* polar lipids (8). This heterogeneity spreads out the filling statistics, and protein occupancy across higher densities of subunit/lipid mole fraction. Further extrusion with smaller 100 & 30 nm filters reduces the rightward tail, leading to a homogeneous distribution of liposomes. The methods of measuring membrane protein reconstitution statistics by proteoliposome efflux assays, co-localization microscopy, and single-molecule photobleaching analysis agree with simulations using the experimental 2:1 POPE:POPG distributions. These results demonstrate that knowledge of the experimentally determined liposome size distribution provides an understanding of the way that protein is captured into vesicles, which can be used to understand when there are conditions where single-molecules prevail, or to extract information about assembly reactions, such as oligomerization (14). In turn, this paper provides key benchmarks for informing on how membrane proteins reconstitute into a synthetic mimic of the *E. coli* membrane, a standard lipid composition for functional assessment of membrane protein structure and function.

At first consideration, it is surprising that EPL and 2:1 POPE/POPG membranes extruded through 400 nm pores generate a population of 25 nm liposomes. However, this has been observed often in liposome populations extruded through filters with pore sizes 200 nm or larger. This effect appears to be independent of the liposome composition, with 1,2-dimyristoyl-sn-glycero-3-phosphocholine (DMPC) (15) and 1,2-dioleoyl-sn-glycero-3-phosphoglycerol (DOPG) (25). Smaller vesicles are also observed in the freeze-fracture micrographs of 400 nm egg PC liposomes, although the overall distribution is described as “relatively homogeneous” (12). In fact, the large standard deviation reported for this distribution indicates that it is likely to contain high polydispersity. In addition, the smaller vesicles are observed across different methods of liposome preparation, including direct hydration of dried lipids, or by solubilization in detergent followed by dialysis. Therefore, it appears as though the presence of small liposomes in the vesicle population may already exist in the sample prior to extrusion, or arise by the physical act of extrusion, independent of the pore size. One way this may occur is by freezing-induced fragmentation. During the freeze-thaw step, smaller liposomes are derived due to the breakage of small membranes that reform into distinct vesicles. This has been demonstrated to occur for dioleoyl-phosphatidylcholine, DOPC membranes (26) as well as mixtures of DOPC and dioleoyl-phosphatidic acid, DOPA (27). In the 2:1 POPE:POPG membrane composition, we observed many smaller vesicles in the freeze-thawed samples, and while we do not know the proportion of these smaller vesicles relative to the true overall distribution, which is not measured here, it suggests that the smaller vesicles are present prior to extrusion. In this case, the sub-population of smaller liposomes acts as a constraint on the heterogeneity of the liposome size distribution. It is an immovable population, and so a homogeneous distribution can only be obtained if the remaining liposomes conform to the contaminating population.

The 30 nm extruded liposome population offers several benefits for the study of membrane proteins in proetoliposomes. It increases the number of liposomes in the sample, and as a result, extends the ‘Poisson-dilution’ range with single or zero occupancy prevailing up to higher densities of 10^-5^ subunits/lipid. This can be useful for the study of transporters, which can have limited signals due to the slow movement of ions or substrates. By extruding samples through 30 nm filters, it is therefore possible to examine the protein at 10-fold higher density while maintaining individual molecules per vesicle. This also offers an ability to maximize the collection of single-molecule microscopy data, for instance dynamic FRET trajectories mapping conformational changes, as one can increase the protein density while increasing the probability that liposomes contain the required single-molecule. Finally, it increases the dynamic range to discriminate between monomer and dimer populations in the freeze-thawed multilamellar membrane, since the chance of random co-capture of subunits is reduced. Our results also indicate that extrusion acts as a passive capture approach and does not affect the characteristics of the system. At low densities, we see no change in the photobleaching probability distributions of CLC-ec1-Cy5 as a function of extrusion pore size. This means that the density dependent monomer-dimer equilirbium in the freeze-thawed large membrane have not been altered due to the act of extrusion alone, supporting extrusion as a passive procedure of compartmentalizing membranes and membrane proteins together.

## 6. Conlcusion

In conclusion, it is important to understand the statistics of membrane protein reconstitution into liposomes that are generally used for studies of membrane protein structure and function. While the commonly used composition of 2:1 POPE:POPG, a mimic of the *E. coli* polar lipids composition, yields an unexpectedly smaller population of liposomes, the heterogeneity in the liposome sizes does not affect the ability to calculate the expected liposome occupancy distribution. Single-molecule microscopy data describing the (i) co-localization fraction of CLC-ec1-Cy5 with AF488-labeled vesicles (ii) and single-molecule photobleaching distributions of CLC-ec1-Cy5, as well as (iii) functional studies of the inactive fraction of vesicles in transport assays of CLC-ec1 or the Fluc F^-^ channel, all agree with the calculated liposome occupancy distribution based on the heterogeneous 400 nm 2:1 POPE:POPG liposome population. Therefore, complete description of liposome occupancy can be caculated for any system provided the liposome size distribution has been measured by cryo-EM or other methods. We recommend proper consideration of this when changing lipid composition in reconstitution studies, as unexpected changes in the size distribution can arise.

## Acknowledgements

We acknowledge Jonathan Remis at the Structural Biology Facility at Northwestern University for help in imaging of liposome samples. The Structural Biology Facility is partially supported by the R.H. Lurie Comprehensive Cancer Center of Northwestern University.

## Funding

This work was supported by the National Institute of General Medical Science, National Institutes of Health [grant numbers R01GM120260, R21GM126476].

## Abbreviations

CHAPS: 3-((3-cholamidopropyl) dimethylammonio)-1-propanesulfonate
DM: n-Decyl-β-D-Maltopyranoside
DOPA: 1,2-dioleoyl-sn-glycero-3-phosphate
DOPC: 1,2-dioleoyl-sn-glycero-3-phosphocholine
POPC: 1-palmitoyl-2-oleoyl-glycero-3-phosphocholine
POPE: 1-palmitoyl-2-oleoyl-sn-glycero-3-phosphoethanolamine
POPG: 1-palmitoyl-2-oleoyl-sn-glycero-3-phospho-(1’-rac-glycerol)
MOPS: 3-(*N*-morpholino)propanesulfonic acid
CLC-ec1: Cl^-^/H^+^ antiporter *E. coli* homologue 1
Cy5: cyanine 5
Cryo-EM: cryogenic electron microscopy
E. coli: *Escherichia coli*
MWCO: molecular weight cut-off
EPL: *E. coli* polar lipids.

