## Supplementary Figure 1 for "Occupancy distributions of membrane proteins in heterogeneous liposome populations"

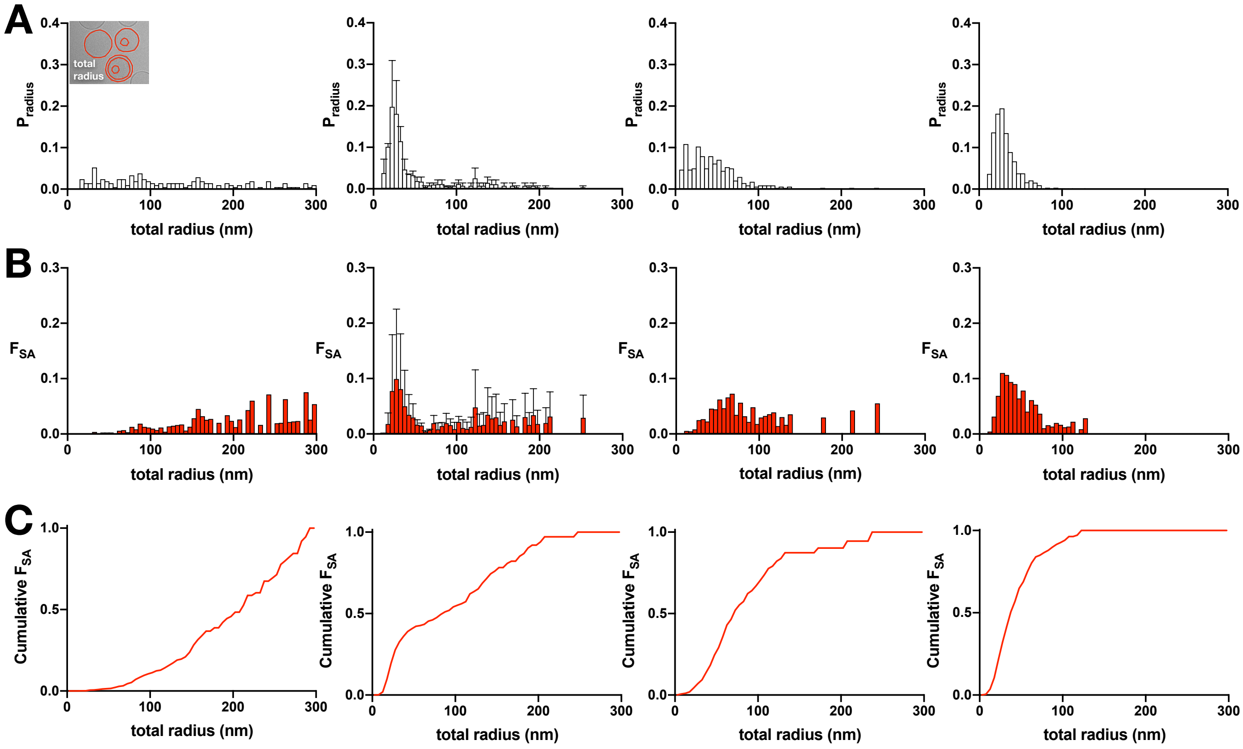


**Supplementary Figure 1. Size distributions of the total radii of extruded liposomes composed of 2:1 POPE:POPG lipids.** (A) Probability distributions of the liposome total radii, $P_{radius}$. (B) Probability distributions of the fractional surface area, $F_{SA}$. (C) Cumulative sum of $F_{SA}$. The data for the 30 nm, 100 nm and 400 nm distributions are presented in Table 2.
