## Supplementary Figure 2 for "Occupancy distributions of membrane proteins in heterogeneous liposome populations"

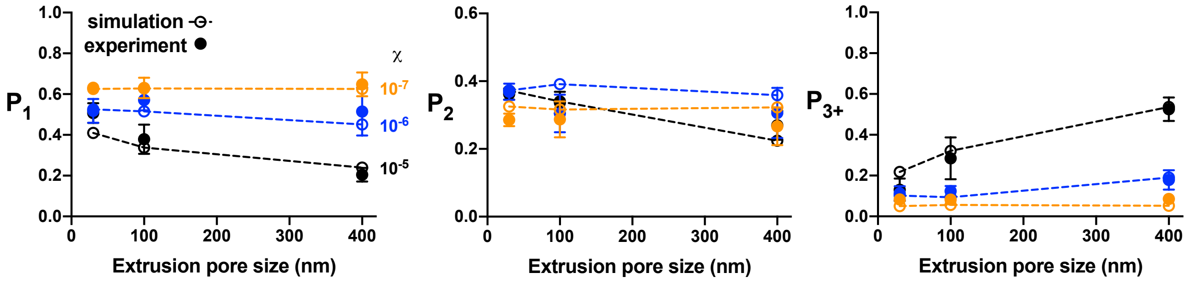


**Supplementary Figure 2. Comparison of experimental and theoretical predictions of photobleaching distributions as a function of liposome extrusion.** Results from Figure 5 plotted to compare theory (hollow circles) vs. experimental data (filled circles).
